# Multimodal imaging for validation and optimization of ion channel-based chemogenetics in nonhuman primates

**DOI:** 10.1101/2023.07.06.547946

**Authors:** Yuki Hori, Yuji Nagai, Yukiko Hori, Kei Oyama, Koki Mimura, Toshiyuki Hirabayashi, Ken-ichi Inoue, Masayuki Fujinaga, Ming-Rong Zhang, Masahiko Takada, Makoto Higuchi, Takafumi Minamimoto

**Author notes:** Authors to whom all correspondence should be addressed: Yuji Nagai, PhD, DVM; Takafumi Minamimoto, PhD. These authors contributed equally to this work (Yuki Hori, Yuji Nagai).

## Abstract

Chemogenetic tools provide an opportunity to manipulate neuronal activity and behavior selectively and repeatedly in nonhuman primates (NHPs) with minimal invasiveness. Designer Receptors Exclusively Activated by Designer Drugs are one example that is based on mutated muscarinic acetylcholine receptors. Another channel-based chemogenetic system available for neuronal modulation in NHPs uses Pharmacologically Selective Actuator Modules (PSAMs), which are selectively activated by Pharmacologically Selective Effector Molecules (PSEMs). To facilitate the use of the PSAM/PSEM system, the selection and dosage of PSEMs should be validated and optimized for NHPs. To this end, we used a multimodal imaging approach. We virally expressed excitatory PSAM (PSAM4-5HT3) in the striatum and the primary motor cortex of two macaque monkeys, and visualized its location through positron emission tomography (PET) with the reporter ligand [^18^F]ASEM. Chemogenetic excitability of neurons triggered by two PSEMs (uPSEM817 and uPSEM792) was evaluated using [^18^F]fluorodeoxyglucose-PET imaging, with uPSEM817 being more efficient than uPSEM792. Pharmacological magnetic resonance imaging showed that increased brain activity in the PSAM4-expressing region began approximately 13 min after uPSEM817 administration and continued for at least 60 min. Our multimodal imaging data provide valuable information regarding the manipulation of neuronal activity using the PSAM/PSEM system in NHPs, facilitating future applications.

**Significance statement:** Like other chemogenetic tools, the ion channel-based system called Pharmacologically Selective Actuator Module/Pharmacologically Selective Effector Molecule (PSAM/PSEM) allows remote manipulation of neuronal activity and behavior in living animals. Nevertheless, its application in non-human primates (NHPs) is still limited. Here, we used multi-tracer positron emission tomography (PET) imaging and pharmacological magnetic resonance imaging (MRI) to visualize an excitatory chemogenetic ion channel (PSAM4-5HT3) and validate its chemometric function in macaque monkeys. Our results provide the optimal agonist, dose, and timing for chemogenetic neuronal manipulation, facilitating the use of the PSAM/PSEM system and expanding the flexibility and reliability of circuit manipulation in NHPs in a variety of situations.

## Introduction

Chemogenetic technology affords remote and reversible control of neuronal activity and behavior in living animals. Chemogenetic tools using mutated muscarinic acetylcholine receptors, called Designer Receptors Exclusively Activated by Designer Drugs (DREADDs), have been widely used in rodents and other small animals (Roth, 2016). DREADDs have also proven to be useful in nonhuman primates (NHPs), where they can manipulate neuronal activity in a specific cell type or neural pathway (Mimura et al., 2021; Oguchi et al., 2021; Perez et al., 2022) or simultaneously and discretely in multiple brain regions (Eldridge et al., 2016; Nagai et al., 2016, 2020; Raper et al., 2019; Hayashi et al., 2020; Hori et al., 2021; Oyama et al., 2021, 2022; Roseboom et al., 2021). As such, they provide an opportunity for identifying the regions and pathways that are causally responsible for cognitive/emotional functions.

Another chemogenetic tool based on engineered ion channels, called Pharmacologically Selective Actuator Module/Pharmacologically Selective Effector Molecule (PSAM/PSEM; Magnus et al., 2011, 2019), has also been developed. In one example, an inhibitory PSAM channel (PSAM4-GlyR) is a chimeric protein made from a modified α7 nicotinic acetylcholine receptor (nAChR) ligand-binding domain that is fused to the anion-permeable ion-pore domain of a glycine receptor. Administering the nicotinic receptor agonist varenicline reduced the firing rate of globus pallidus neurons in a monkey only after transduction of PSAM4-GlyR (Magnus et al., 2019). The same study showed that two selective and potent PSEMs, uPSEM817 and uPSEM792, effectively modulated mouse behavior via PSAM4-GlyR. Additionally, these two PSEMs showed good brain penetrance, relatively slow washout, negligible conversion to potential metabolites, and no significant effects on heart rate or sleep in monkeys, suggesting that they are suitable candidates for NHP studies (Raper et al., 2022). The PSAM/PSEM system has advantages over DREADDs, particularly the ability to directly modulate neuronal electrical activity. When used with DREADDs in the same monkey, it allows for complex and flexible chemogenetic control, including multiplexed manipulations, increasing the utility of NHPs as neurobiological models. However, before widespread use in NHP studies is possible, several factors should be validated and optimized. Specifically, to reduce the burden on internal organs such as the liver, minimizing the necessary total amount of activator/solvent is crucial. Therefore, using activators that induce chemogenetic action at low concentrations is preferable.

For the application and optimization of DREADDs in NHPs, the use of non-invasive imaging, such as positron emission tomography (PET), has proven to be valuable. PSAM4 visualization via PET imaging with the radiolabeled α7 nAChR antagonist [^18^F]ASEM (Horti et al., 2014) has been demonstrated in rats (Magnus et al., 2019), suggesting that it may be useful for verification of PSAM4 gene transduction in NHPs. In addition, PET with [^18^F]fluorodeoxyglucose (FDG) has been used to assess chemogenetic functions, allowing the agonist dose-response relationship to be determined (Bonaventura et al., 2019; Nagai et al., 2020). Moreover, pharmacological magnetic resonance imaging (phMRI) has been shown to be a powerful tool for measuring dynamic blood oxygen-level dependent (BOLD) signal changes induced by administration of DREADD ligands (Roelofs et al., 2017; Peeters et al., 2020), suggesting that it could also be used to reveal the functional dynamics of the PSAM/PSEM system.

Here, we aimed to validate and optimize the PSAM/PSEM system for macaque monkeys using a multimodal imaging strategy (Fig. 1). We virally expressed the PSAM4-5HT3 (excitatory; PSAM4 combined with serotonin receptor 3) (Magnus et al., 2019) in the striatum and the primary motor cortex (MI) (Fig. 1A). We then performed a multitracer ([^18^F]ASEM and [^18^F]FDG) PET study to evaluate the efficacy of two PSEMs, uPSEM817 and uPSEM792. [^18^F]ASEM was used to visualize expression (Fig. 1B) and [^18^F]FDG was used to assess chemogenetic excitability (Fig. 1C). We further investigated the time-course of chemogenetically-induced BOLD signal changes using phMRI (Fig. 1D). Our multimodal imaging data provide useful information about the PSAM/PSEM system for manipulating neuronal activity in NHPs.

**Figure 1.**
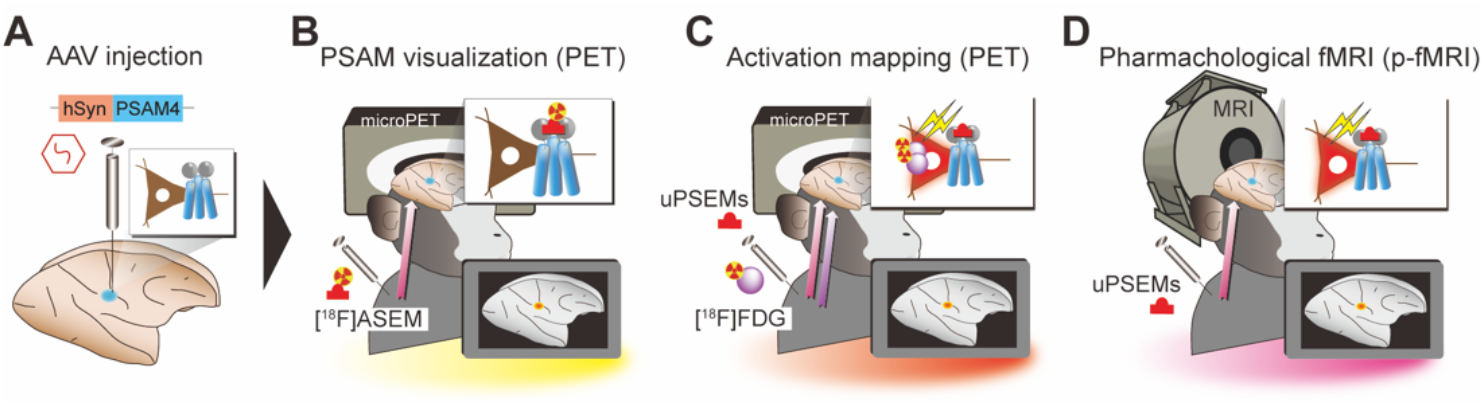
Experimental design. **A**. Monkeys received AAV-PSAM4 vector injections into the striatum (n = 2 monkeys) and primary motor cortex (MI, n = 1). **B**. [^18^F]ASEM-PET was performed to visualize PSAM4 expression *in vivo*. **C**. After administering uPSEMs, activation mapping using [^18^F]FDG-PET imaging allowed the relationship between agonist type/dose and activation strength to be determined. **D**. Pharmacological MRI was used to measure the time-course of chemogenetically induced activity changes.

## Materials and Methods

### Animals

Two macaque monkeys participated in the experiments (MK#1: Cynomolgus monkey [*Macaca fasciculus*]; male, 5.9 kg, aged 7 years at the start of experiments; MK#2: Japanese monkey [*Macaca fuscata*]; male, 7.3 kg, aged 9 years at the start of experiments) (Table 1). All experimental procedures involving animals were carried out in accordance with the Guide for the Care and Use of Nonhuman primates in Neuroscience Research (The Japan Neuroscience Society; https://www.jnss.org/en/animal_primates), were approved by the Animal Ethics Committee of the National Institutes for Quantum Science and Technology, and were consistent with the ARRIVE (Animal Research: Reporting of In Vivo Experiments) guidelines. A standard diet, supplementary fruits/vegetables, and a vitamin C tablet (200 mg) were provided daily. *C*, cynomolgus monkey; *J*, Japanese monkey; *PUT*, putamen; *LC*, low conductance; *CDh*, head of the caudate nucleus; *CDt*, caudate tail; *MI*, primary motor cortex. Check marks indicate that the experiments were performed.

**Table 1.**
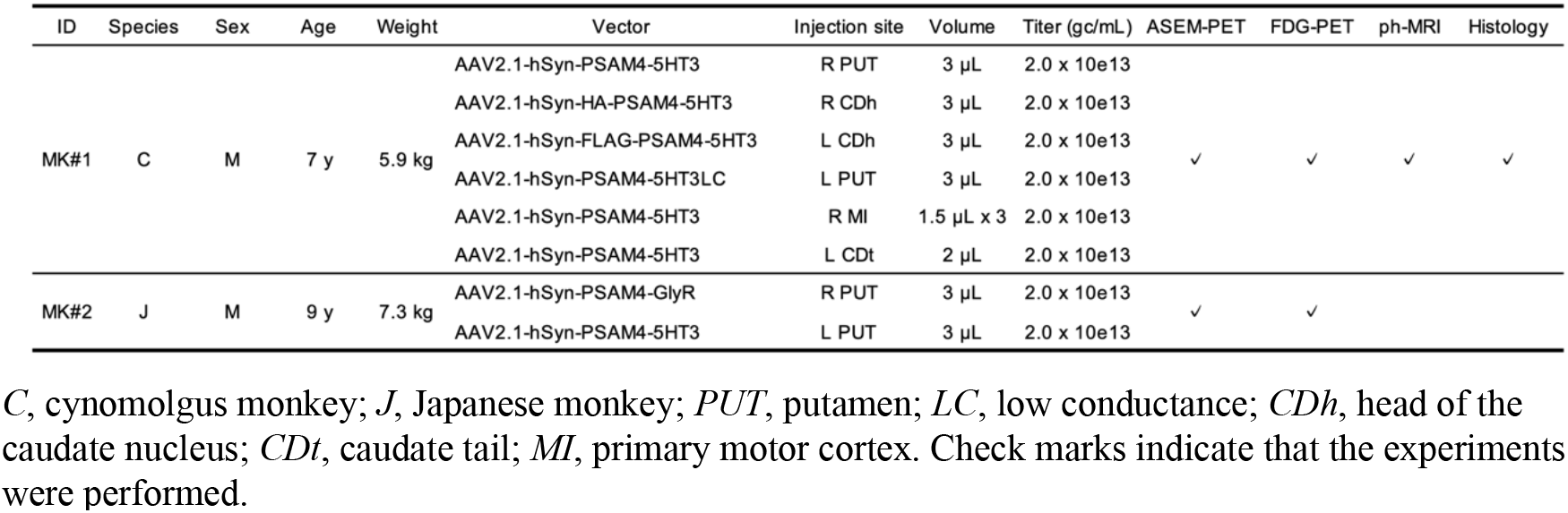
Summary of vector injection and experiments for each monkey

### Viral vector production

Viral vectors used in this study and their titers are summarized in Table 1. The adeno-associated virus (AAV) vectors were produced by helper-free triple transfection and purified using affinity chromatography (GE Healthcare, Chicago, IL, USA). For production of AAV2.1 vector, the pAAV-RC1 plasmid-coding AAV1 capsid protein and the pAAV-RC2 plasmid-coding AAV2 capsid protein were transfected at a 1:9 ratio. Viral titer was determined by quantitative PCR using Taq-Man technology (Life Technologies, Waltham, MA, USA). Additional details have been described previously (Kimura et al., 2023). cDNA fragments encoding PSAM4, such as PSAM4-5HT3, were synthesized based on the sequences originally reported by Magnus et al. (2019). We always used a high conductance variant of PSAM4-5HT3, except for a single case in which we used a low conductance variant (PSAM4-5HT-LC). The transfer plasmid was constructed by replacing the CMV promoter of an AAV backbone plasmid (pAAV-CMV, Stratagene, San Diego, CA, USA) with a human Syn promoter, and then inserting the cDNA fragment and the WPRE sequence.

### Surgical procedures and viral vector injections

MK#1 received the first injections of viral vectors into the bilateral head of the caudate nucleus (CDh) and putamen (PUT). Six months later, the second vector injections were performed in the right primary motor cortex (MI) and the left caudate tail (CDt). MK#2 received vector injections into the bilateral PUT. AAV vectors and injection sites are summarized in Table 1. Surgeries were performed under aseptic conditions in a fully equipped operating suite. We monitored body temperature, heart rate, SpO_2_, and tidal CO_2_ throughout all surgical procedures. Anesthesia was induced using intramuscular (i.m.) injection of ketamine (5–10 mg/kg) and xylazine (0.2–0.5 mg/kg), and monkeys were intubated with an endotracheal tube. Anesthesia was maintained with isoflurane (1%–3%, to effect). After surgery, prophylactic antibiotics and analgesics were administered. Before surgery, X-ray computed tomography (CT, 3D Accuitomo 170, J. Morita, Kyoto, Japan) and 7 Tesla magnetic resonance (MR) images (BioSpec 70/40, Bruker) were acquired under anesthesia (continuous infusion of propofol 0.2–0.6 mg/kg/min, intravenously). Overlaid MR and CT images were created using PMOD® image analysis software (PMOD Technologies Ltd., Zurich, Switzerland) to estimate the stereotaxic coordinates of target brain structures.

For the striatum, the monkeys received injections with AAV vectors as described in a previous study (Nagai et al., 2016). Briefly, viruses were pressure-injected using a 10-μL microsyringe (Model 1701RN, Hamilton) with a 30-gauge injection needle placed in a fused silica capillary (450 mm OD), which minimizes backflow by creating a 500-μm space surrounding the needle tip. The microsyringe was mounted into a motorized microinjector (UMP3T-2, WPI) that was held by a manipulator (Model 1460, David Kopf, Ltd.) on the stereotaxic frame. After a burr hole (8-mm diameter) and the dura mater (∼ 5 mm) were opened, the injection needle was inserted into the brain and slowly moved down 2 mm beyond the target and then kept stationary for 5 min, after which it was pulled up to the target location. The injection speed was set at 0.2–0.5 μL/min. After the injection, the needle remained *in situ* for 15 min to minimize backflow along the needle.

MI injections in MK#1 were performed under direct vision using the same type of surgical procedures as described previously (Miyakawa et al., 2023). The putative hand/arm target region of the right MI was determined based on sulcal landmarks from a pre-surgical MRI. After retracting the skin and galea, the cortex was exposed by removing a bone flap and reflecting the dura matter ventrally. Vectors were pressure-injected using a 10-μL syringe with a 33-gauge needle (NanoFil Syringe, WPI, Sarasota, FL, USA). The syringe was mounted on a motorized microinjector (UMP3T-2, WPI) that was attached to a manipulator (Model 1460, David Kopf, Ltd., Tujunga, CA, USA) on the stereotaxic frame. For each injection, the injection needle was inserted into the brain and moved down to 3.0 mm below the surface. After 5 min, the needle was pulled up 0.5 mm, and 1.5 μL of the vector solution was injected at 0.15 μL/min. The needle was then slowly pulled up following an additional 5-min waiting period to prevent backflow. In total, virus vector injections were conducted at three sites (one depth per site) within the target region, with an inter-site distance of approximately 1.2 to 1.5 mm along the cortical surface (Fig. 1C). The dura matter was sutured back with absorbable material (VICRYL Plus, Johnson & Johnson, New Brunswick, NJ, USA) and the bone flap was also replaced and fixed to the skull with absorbable suture via drilled cranial holes before closing the galea and the skin.

### Drug preparation and administration

We used uPSEM817 (uPSEM817 tartrate, TOCRIS #6866), uPSEM792 (uPSEM792 hydrochloride, TOCRIS #6865), and varenicline (varenicline tartrate, TOCRIS #3754) as PSAM4 ligands (Magnus et al., 2019). uPSEM817 and uPSEM792 were dissolved in 0.9% saline, while varenicline was dissolved in 2% dimethyl sulfoxide (DMSO, FUJIFILM Wako Pure Chemical) and diluted in 0.9% saline. The ligand doses ranged from 1 to 300 μg/kg and were not corrected for the counterion mass. All drugs were administrated intravenously (i.v.).

### PET imaging

PET imaging was conducted to examine the expression of PSAM4 *in vivo* with [^18^F]ASEM as the PET ligand. Briefly, the monkey was anesthetized with ketamine (5–10 mg/kg, i.m.) and xylazine (0.2–0.5 mg/kg, i.m.), and kept anesthetized with isoflurane (1%–3%) during all PET procedures. PET scans were performed using a microPET Focus 220 scanner (Siemens Medical Solutions, Malvern, PA, USA). Transmission scans were performed for about 20 min with a ^68^Ge source.

Emission scans were acquired in 3D list mode with an energy window of 350–750 keV after an intravenous bolus injection of [^18^F]ASEM (198.2–263.9 MBq). Emission-data acquisition lasted 90 min. PET imaging was performed before vector injection (i.e., baseline scan) and about five to six weeks after the injection when expression was expected to be established, as previously reported (Nagai et al., 2016, 2020).

To examine the effect of chemogenetic activation, PET scan with [^18^F]FDG (injected radioactivity, 183.5–226.1 MBq) was performed. PET procedures were the same as for [^18^F]ASEM, except that emission scans started 1 min after administration of the PSEMs or vehicle.

PET images were reconstructed with filtered back-projection using a Hanning filter cutoff at a Nyquist frequency of 0.5 mm^−1^. Dynamic PET images were converted to standardized uptake values (SUV, regional radioactivity [Bq cm^−3^] × body weight (g)/injected radioactivity [Bq]). PET SUV images were averaged between 60 and 90 min for [^18^F]ASEM and between 30 and 60 min for [^18^F]FDG, and SUV ratio (SUVR) images were created by normalizing to the mean value of the whole brain. The expression of PSAM4 was estimated by the change in SUVR (ΔSUVR; post-injection value – pre-injection value). Chemogenetically induced metabolic changes were estimated by the ΔSUVR of [^18^F]FDG, i.e., the pretreatment value for vehicle subtracted from the pretreatment value for the PSEMs. Subtraction images were constructed using SPM12 software (The Wellcome Centre for Human Neuroimaging; www.fil.ion.ucl.ac.uk) and MATLAB R2016a software (MathWorks Inc., Natick, MA, USA). To quantify the metabolic changes, volumes of interest (VOIs) were defined for each PSAM4-positive region in which the baseline subtracted value was higher than half of the highest value for each vector injection site. A Gaussian filter with a 2-mm full-width-at-half-maximum was applied to the ΔSUVR FDG images.

### Pharmacological MRI

To assess the dynamic changes in chemogenetically induced BOLD signals, phMRI was performed for MK#1 using a custom made 8-channel receiver coil (Takashima Co. Ltd., Tokyo, Japan) and a 7 Tesla MRI (BioSpec 70/40, Bruker, Ettlingen, Germany). The monkeys were anesthetized with ketamine (5–10 mg/kg, i.m.) and xylazine (0.2–0.5 mg/kg, i.m.), followed by step-down infusion of propofol (0.2-0.6 mg/kg/min i.v.), as in a previous report (Miyabe-Nishiwaki et al., 2013). PhMRI data were acquired two times for each condition (uPSEM817 or vehicle) using gradient-echo multi-slice sequences, as in a previous study (Roelofs et al., 2017). Acquisition parameters were as follows; TR / TE = 1250 / 5 ms, flip angle = 60°, field of view = 84 × 84 × 48 mm, matrix size = 70 × 70 × 32, and total scan time was 1 min per 1 volume. After 5 or 15 baseline images were acquired, vehicle or uPSEM817 (300 μg/kg) was injected via the crural vein and followed by acquisition of another 40 to 60 images. PhMR images were corrected for motion using FLIRT; the brain images were extracted and normalized to the Yerkes19 macaque template (Donahue et al., 2016) using BET, FLIRT, and FNIRT, and the values in all images were scaled to those in mean baseline images in each scan at the voxel level. All tools were implemented in FSL software (FMRIB’s Software Library, http://www.fmrib.ox.ac.uk/fsl) (Smith et al., 2004).

### Histology and immunostaining

For histological examination, MK#1 was immobilized with ketamine (10 mg/kg, i.m.) and xylazine (0.5 mg/kg, i.m.), deeply anesthetized with an overdose of thiopental sodium (50 mg/kg, i.v.), and then transcardially perfused with saline at 4°C, followed by 4% paraformaldehyde in 0.1 M phosphate-buffered saline (PBS) at a pH of 7.4. The brain was removed from the skull, postfixed in the same fresh fixative overnight, saturated with 30% sucrose in phosphate buffer at 4°C, and then cut serially into 50-μm-thick sections on a freezing microtome. For visualization of immunoreactive signals against FLAG-tag and HA-tag, every 4th section was immersed in 1% skim milk for 1 h at room temperature and incubated overnight at 4°C with mouse anti-FLAG monoclonal antibody (1:500; MAB3118, Sigma-Aldrich) or mouse anti-HA monoclonal antibody (1:500; 901513, Biolegend), and then for 2 days in PBS containing 0.1% Triton X-100 and 1% normal goat serum at 4°C. The sections were then incubated in the same fresh medium containing biotinylated goat anti-mouse IgG antibody (1:1,000; 715-065-150, Jackson Immuno Research) for 2 h at room temperature, followed by avidin-biotin-peroxidase complex (ABC Elite, Vector Laboratories) for 2 h at room temperature. For visualization of the antigen with diaminobenzidine (DAB), the sections were reacted in 0.05 M Tris-HCl buffer (pH 7.6) containing 0.04% DAB, 0.04% NiCl_2_, and 0.003% H_2_O_2_. Then, the sections were nucleus-stained with 0.5% Neutral red, mounted on gelatin-coated glass slides, air-dried, and cover-slipped. To achieve double immunofluorescence immunohistochemistry for FLAG and NeuN, sections were pretreated as described above, and then incubated with mouse monoclonal anti-FLAG antibody (1:500; MAB3118, Sigma-Aldrich) and anti-NeuN antibody (1:1,000, MABN140, Sigma-Aldrich) for 2 days at 4 °C. The sections were subsequently incubated for 2 h at room temperature with a cocktail of Alexa 488-conjugated donkey anti-rabbit IgG antibody (1:400 dilution; A-11008, Invitrogen) and Alexa 568-conjugated donkey anti-mouse IgG antibody (1:400 dilution; A-11004, Invitrogen). Bright-field images of sections were captured using a digital slide scanner (NanoZoomer S60, Hamamatsu Photonics K.K., Hamamatsu, Japan; 20× objective, 0.46 μm per pixel). Florescence images of sections were digitally captured using an optical microscope equipped with a high-grade charge-coupled device (CCD) camera (Biorevo, Keyence, Osaka, Japan; 20× objective). JPEG images were exported using a viewer software (NDP.view2, Hamamatsu Photonics K.K.; BZ-X Analyzer, Keyence).

### Statistical Analysis

All statistical analyses were performed in the R statistical computing environment (R Development Core Team, 2004). A one-way repeated-measures analysis of variance (ANOVA) was performed to assess whether the FDG uptake depended on the concentration of uPSEM817. Comparison of FDG uptake between the two PSEMs at a dose of 100 μg/kg was tested by two-tailed paired t-test. In the phMRI experiment, BOLD signal changes at each PET-defined VOI were measured repeatedly at 1-min intervals in two independent sessions. These two datasets were then averaged and treated as the time course data. BOLD signal changes before (−15–0 min) and after (20–35 min) administration were compared using Wilcoxon rank-sum test with post-hoc Bonferroni correction. To assess the dynamics of the BOLD signal changes in the PSAM4-expressing regions, the time course of the BOLD signal was plotted and then fitted to a sigmoid curve with three parameters according to the following equation:

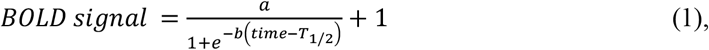

where *a* is the BOLD signal at the plateau, *b* is the time constant, and *T*_*1/2*_ is the time halfway between initial rise and the start of the plateau. The data and code used to generate the results are available in the Open Science Framework database (https://github.com/minamimoto-lab/2023-Hori-PSAMPSEM).

## Results

### PET imaging validation of virally expressed PSAM4

MK#1 received injections of four different AAV vectors (PSAM4-5HT3, PSAM4-5HT3-LC, HA-PSAM4-5HT3, and FLAG-PSAM4-5HT3) for expressing PSAM4-5HT3 under the drive of the human synapsin promoter (hSyn), one for each of four striatal locations (CDh and PUT in each hemisphere; Fig. 2A; Table 1). We performed PET imaging with [^18^F]ASEM, a radiotracer for PSAM4 (Magnus et al., 2019), before the injection and 6 weeks afterward. Figure 2B shows coronal sections of the parametric PET image showing regional [^18^F]ASEM uptake relative to the whole brain. Baseline images before AAV transduction showed weak uptake of [^18^F]ASEM in the anterior cingulate region, likely reflecting tracer binding to endogenous α7 nAChR, for which [^18^F]ASEM also has strong affinity (Fig. 2B, left). The post-injection image showed regions with enhanced uptake in the CDh and PUT, which corresponded to the AAV injection sites. The subtraction images showed increased uptake in the right and left PUT, corresponding to the injection sites of the AAV vectors for PSAM4-5HT3 and PSAM4-5HT3-LC expression, respectively (Fig. 2C, left). A moderate increase in uptake was also observed in the right and left CDh, corresponding to the injection sites of the AAV vectors for expressing HA-PSAM4-5HT3 and FLAG-PSAM4-5HT3, respectively (Fig. 2C, left). Post-mortem immunohistology revealed positive immunolabeling with anti-FLAG antibody in the left CDh (corresponding to the FLAG-PSAM4-vector injection sites; Fig. 2D), suggesting that our PET results reflect *in vivo* detection of PSAM4 expression. Double immunofluorescence staining further confirmed PSAM4 expression in neurons (Fig. 2E). In addition, we also found increased uptake in the bilateral substantia nigra, corresponding to the striatal projection sites, presumably reflecting PSAM4-5HT3 expression at the axon terminals. Positive signals could not be detected by immunohistological analysis using anti-HA antibodies (see Discussion).

**Figure 2.**
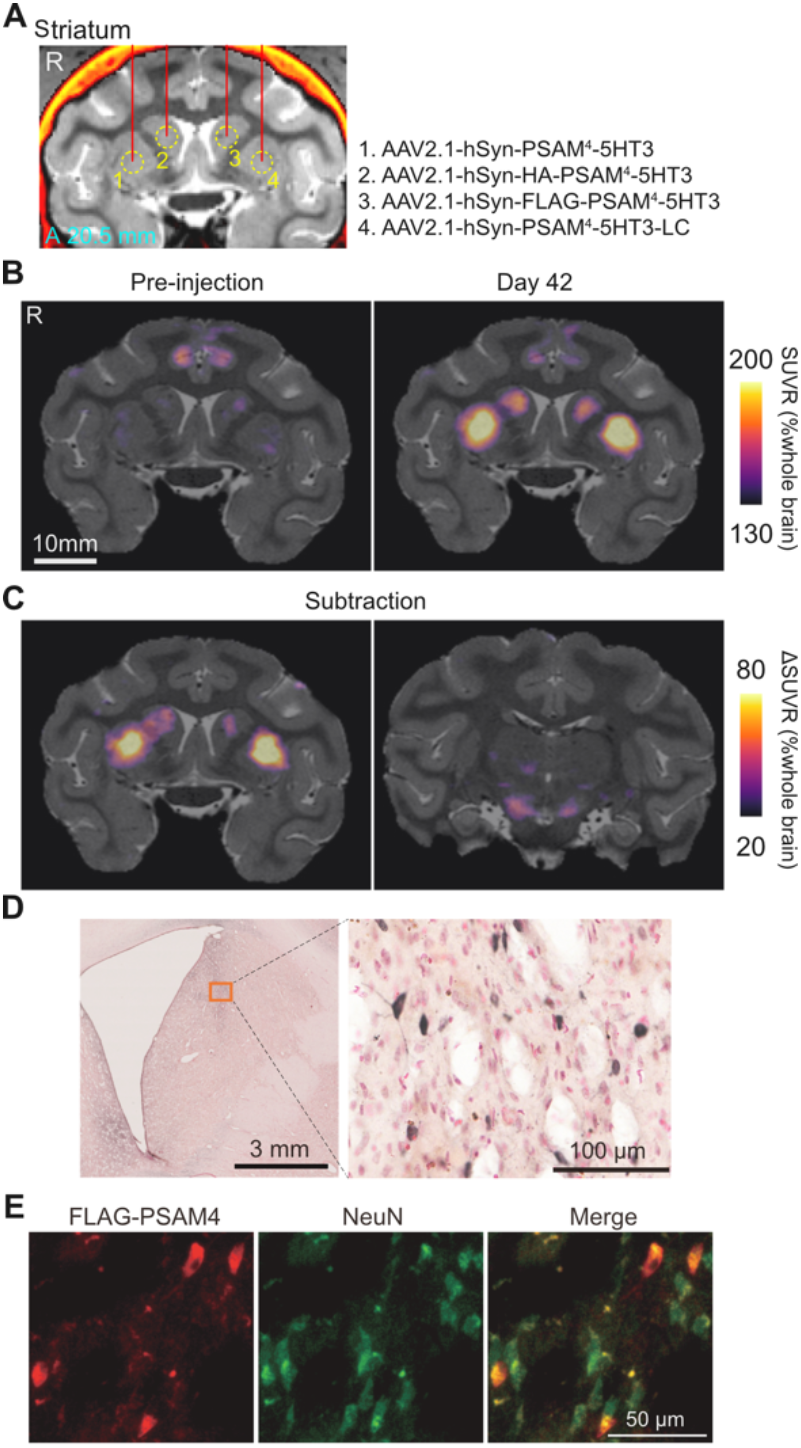
PET imaging visualizing PSAM4-5HT3 expression *in vivo*. **A**. Location of viral vector injections on a coronal CT-MR fusion image, 20.5 mm anterior to the ear-bar zero line. AAV2.1 vectors expressing PSAM4-5HT3 were injected into the striatum. The viral vector constructs and the injected regions correspond by number. **B**. Coronal PET-MR fusion images before and 42 days after vector injection. PET signals are expressed as SUVR, which was normalized by whole-brain SUV and generated by averaging the dynamic PET data 60 to 90 min after i.v. injection of [^18^F]ASEM. **C**. Contrast (subtraction) of SUVR images taken before and 42 days after vector injection. Increased SUVR was observed in the striatum where the vector was injected (left) and in the substantia nigra, to which the striatal neuron projected (right). **D**. Histological confirmation of PSAM4-5HT expression. Microscopic image of a DAB immunostained section of anti-FLAG positive neurons in the left caudate nucleus, where an hSyn-FLAG-PSAM4 vector was injected. **E**. Double immunofluorescence staining of anti-FLAG (red) and anti-NeuN (green). Data obtained from MK#1.

We then performed another set of AAV injections (AAV2.1-hSyn-PSAM4-5HT3) in the right MI and left CDt. The PET scans with [^18^F]ASEM showed a substantial increase in tracer uptake in the PSAM4-vector-injected side of the MI (Fig. 3A, red arrow). In addition, uptake increased modestly in the ipsilateral thalamus, PUT, and contralateral MI, which appears to reflect expression in the axon terminals of neurons in the target MI (Fig. 3B, yellow, white and red triangles). A weak but noticeable increase in uptake was also observed in the left CDt (Fig. 3B, left, yellow arrow). Collectively, these results suggest that our AAV vector system effectively delivers PSAM4-5HT3 in monkeys, which can be visualized *in vivo* by [^18^F]ASEM-PET.

**Figure 3.**
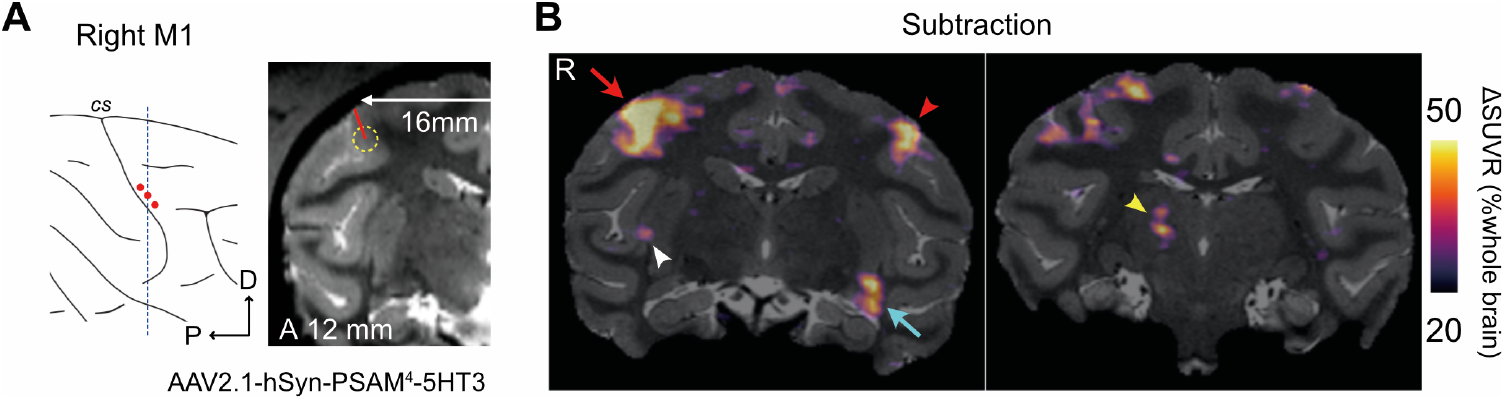
PET imaging visualizing PSAM4-5HT3 expression in the cortex and its projection sites. **A**. Illustration and MR image of the location of the second vector injection site. The vector was injected in three tracks within the right MI, shown as red dots in the illustration. **B**. The SUVR images contrasting 42 days after the second and first vector injection. Increased SUVR was observed in the right MI (left, red arrow) and left CDt (left, cyan arrow). Other projection sites, such as the left MI (left, red arrowhead), right PUT (left, white arrowhead), and right thalamus (right, yellow arrowhead), also showed increased SUVR. Data obtained from MK#1.

### PET assessment of chemogenetic activation induced by agonists

Next, we performed a PET study with [^18^F]FDG to assess whether administration of the chemogenetic ligands (the PSEMs) activated PSAM4-5HT3 and increased regional brain glucose metabolism, as an index of neuronal/synaptic activation in the brain (Nagai et al., 2020). MK#1 underwent [^18^F]FDG-PET imaging following intravenous administration of the two PSEMs (uPSEM817 or uPSEM792), varenicline (another less-selective agonist), or vehicle as a control. Compared with that following administration of vehicle control, FDG uptake in the PSAM4-5HT3-positive striatal regions increased significantly in a dose-dependent manner following administration of uPSEM817 (repeated-measures one-way ANOVA; *F*_[1, 11]_ = 31.97; *p* < 0.001) (Fig. 4A, 4B, left). When administered at 100 μg/kg, uPSEM817 and varenicline increased FDG uptake by an average of 6.4% and 3.6%, respectively, while uPSEM792 showed a limited increase in uptake (0.7%) (Fig. 4B). Comparing the two PSEMs, FDG uptake after administration of uPSEM817 was higher than it was for uPSEM792, although this was not statistically significant (Wilcoxon signed rank exact test; *p* = 0.125). An increase in FDG uptake (18.5%) was also observed in the PSAM4-5HT3 positive MI regions following uPSEM817 administration (100 μg/kg) (Fig. 4C).

**Figure 4.**
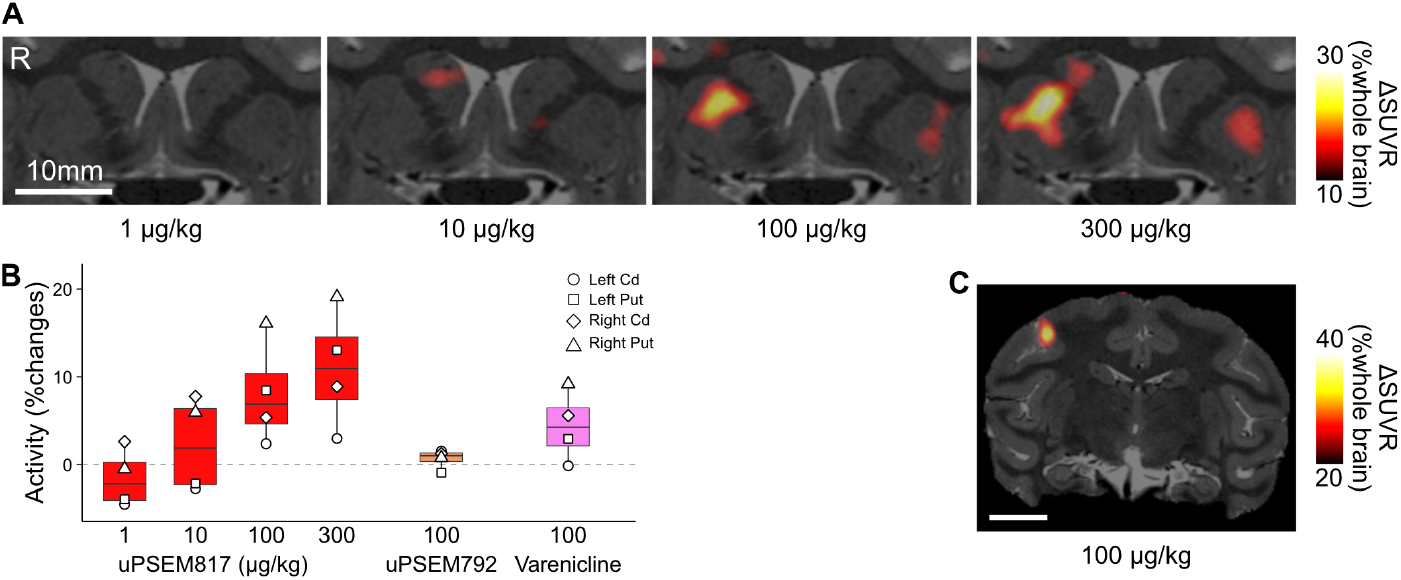
FDG-PET assessment of chemogenetic activation using PSEMs. **A**. Coronal sections of ΔSUVR [^18^F]FDG-PET images overlayed with an MR image indicate chemogenetically induced metabolic changes via PSAM4-5HT3 activation in striatal regions following uPSEM817 administration (1 to 300 μg/kg, n = 1 for each dose). ΔSUVR (color coded) indicates the change in SUVR (values for uPSEM817 minus those for vehicle). **B**. Comparison of chemogenetic effects of different ligands (red: uPSEM817; orange: uPSEM792; pink: varenicline) and their doses. Chemogenetically induced metabolic activity (bars and symbols indicate averages and striatal VOIs, respectively) was normalized by the values from the vehicle condition and plotted. **C**. A coronal SUVR contrast image overlayed with an MR image expressing PSAM4-5HT3 in the MI. Data obtained from MK#1.

### Across-animal reproducibility of the PET imaging-based validation

We next validated our PET imaging-based assessment of PSAM4 expression and function in another monkey (MK#2). The left PUT of MK#2 was injected with an AAV vector expressing excitatory PSAM4 and the right PUT was injected with one expressing inhibitory PSAM4 (Table1; Fig. 5A). Five weeks later, [^18^F]ASEM-PET visualization indicated increased uptake in the left PUT (PSAM4-5HT3-side), while the uptake in the right PUT was unchanged (PSAM4-GlyR-side) (Fig. 5B). Although [^18^F]ASEM-PET failed to visualize PSAM4-GlyR expression, it is a reproducible and useful tool for at least detecting PSAM4-5HT3-expressing regions *in vivo*.

**Figure 5.**
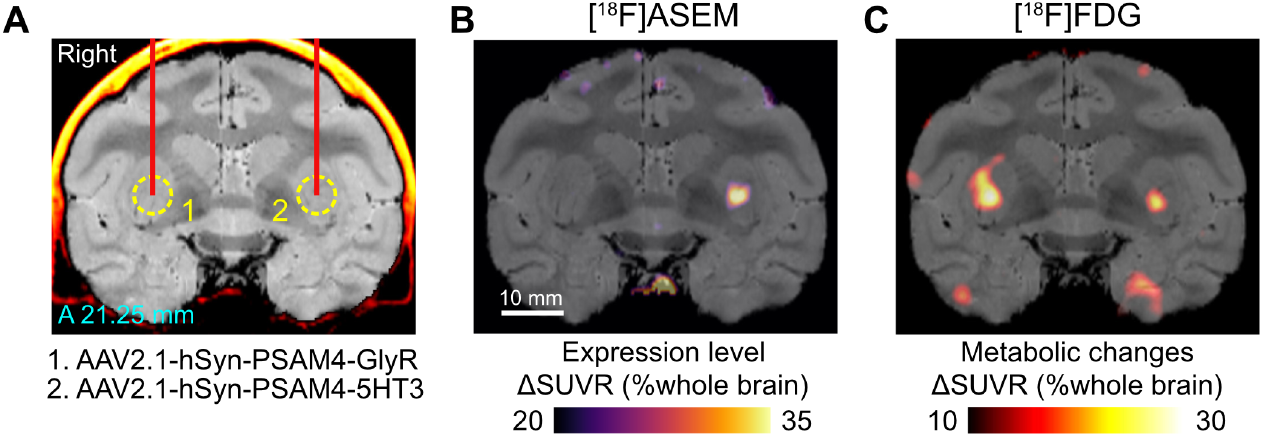
PET validation of PSAM4 expression and function in MK#2. **A**. Location of viral vector injections are shown on a coronal CT-MR fusion image, 21.25 mm anterior to the ear-bar zero line. AAV2.1 vectors expressing PSAM4-GlyR and PSAM4-5HT3 were injected into the right and left putamen, respectively. **B**. ΔSUVR image showing increased [^18^F]ASEM uptake in the striatum where the vector expressing PSAM4-5HT3 was injected. Notable increase was not found in the right PUT (PSAM4-GlyR-side). **C**. A coronal ΔSUVR section of [^18^F]FDG-PET overlayed with an MR image indicate increased metabolic activity following uPSEM817 administration (300 μg/kg, iv). Data obtained from MK#2.

We next sought to examine the effect of chemogenetic activation. Consistent with the results from MK#1, [^18^F]FDG-PET demonstrated increased metabolic activity in the PSAM4-5HT3-expression site following administration of uPSEM817 (300 μg/kg, iv) (Fig. 5C). Counterintuitively, in the right PUT, where an AAV vector-expressing inhibitory PSAM (PSAM4-GlyR) was injected, the metabolic activity was also 10% greater than in the control (Fig. 5C). This probably reflects depolarization of striatal projection neurons via activation of PSAM4-GlyR, which has been demonstrated in rodent slice preparations (Gantz et al., 2021). Collectively, these results demonstrate the across-animal reproducibility of PET visualization and validation of PSAM4 expression/function.

### Pharmacological MRI assessment of chemogenetically induced activity changes

Next, the time course of chemogenetically induced activity change was assessed using phMRI (Roelofs et al., 2017). Compared with those during the pre-injection baseline period, after uPSEM817 administration (300 μg/kg, iv) in MK#1, the BOLD signals increased significantly in the right MI, right PUT, and right CDh, which corresponded to the PSAM4 expressing regions as indicated by [^18^F]ASEM-PET (*p* < 0.001, Wilcoxon rank-sum test with post-hoc Bonferroni correction for multiple comparisons; Fig. 6A), confirming the FDG-PET results obtained in the same monkey (Fig. 4). A similar but non-significant increase in the BOLD signal was induced in the left CDh and PUT. The BOLD signal was unchanged in the brain regions that did not express PSAM4, such as insular cortex (Fig. 6A, R INS). Control vehicle administration did not induce a significant BOLD change in any of these brain regions (*p* > 0.05, data not shown but see Fig. 6B-D, cyan). To determine the timing of the BOLD signal increase associated with chemogenetic neuronal activation, sigmoid curves with three parameters were fitted to the BOLD data (Fig. 6B-D). The intermediate time between the initial increase and the beginning of the plateau (*T*_*1/2*_) was roughly consistent among the three regions at approximately 13 min (12.9, 13.1, and 13.8 min for right MI, right PUT, and right CDh, respectively). Moreover, the increase in BOLD signal was maintained in each of the three regions for at least one hour after uPSEM817 administration.

**Figure 6.**
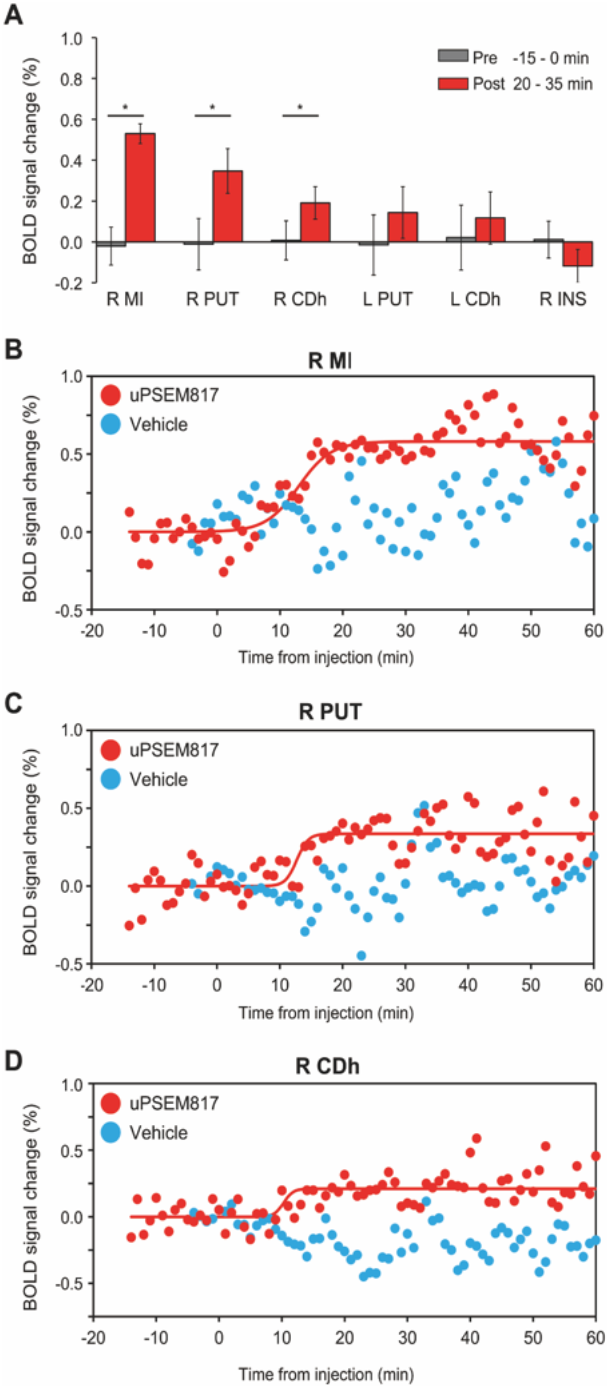
Pharmacological MRI assessment of chemogenetically induced activity changes in PSAM4-5HT3 expressing regions. **A**. Comparison of the BOLD signal changes between pre-injection (0 to 15 min before injection: grey bar) and post-uPSEM817 injection (20 to 35 min after injection: red bar) periods for each region in which PSAM4 was expressed and for a control region in which it was not expressed (right insula, R INS). The red bars show the BOLD signal changes over 15 min (i.e., fifteen time points; error bars are the standard deviation). Asterisks indicate the statistical significance between pre-and post-injection differences (*p* < 0.001, Wilcoxon rank-sum test with post-hoc Bonferroni correction for multiple comparisons). Note that activity did not differ significantly between pre- and post-vehicle injection (*p* > 0.05). **B-D**. Time courses of BOLD signal changes before and after uPSEM817 (red dots) or vehicle injections (cyan dots) in the R MI (B), R PUT (C), and R CDh (D). Red curves represent best-fit sigmoid functions to the data (Eq. 1). Parameter *a*, the BOLD signal increase when signals reach plateau, was equivalent to the results shown in **A**; 0.58%, 0.34%, and 0.21% for the R MI, R PUT, and R CDh, respectively. The time halfway between initial rise and the start of the plateau (*T*_*1/2*_) was 12.9 min, 13.1 min, and 13.8 min for the R MI, R PUT, and R CDh, respectively. Data obtained from MK#1 in two independent sessions. L, left; R, right; MI, primary motor cortex; PUT, putamen; CDh, head of the caudate nucleus.

## Discussion

In the present study, we applied imaging-based strategies to optimize and validate the function of the PSAM/PSEM chemogenetic system in NHPs. PET imaging with a PSAM probe, [^18^F]ASEM, was used to verify PSAM4 gene transduction and to estimate the location and level of expression, and [^18^F]FDG-PET was used to compare metabolic changes after administration of the two PSEMs. PhMRI was used to measure the time course of chemogenetically induced changes in BOLD activity. Taken together, we provide important data for optimizing agonist, dosage, and timing of chemogenetic neuronal manipulation using PSAM4 in monkeys. We hope these optimized values will help increase the application of these tools in NHPs for a variety of applications.

### Multimodal imaging assists in verification of gene transfer and optimization of actuators

In the current study, we used AAV2.1 vectors with a human synapsin promoter (hSyn) for neuron-specific transduction of PSAM4-5HT3 with or without fusion protein tags. The combination of AAV2.1, hSyn, and HA/FLAG tags has been used in this study and in previous hM4Di/hM3Dq DREADD studies with macaque monkeys (Kimura et al., 2023; Miyakawa et al., 2023). As expected, our AAV construct successfully induced PSAM4-5HT3 expression in the striatum and MI cortex, as visualized by [^18^F]ASEM-PET. In contrast to untagged PSAM4 vectors, using PSAM4 vectors with fusion tags (HA and FLAG) resulted in relatively weaker [^18^F]ASEM signal and FDG uptake, which is consistent with previous observations that AAV constructs with a fusion of protein tags led to decreased expression levels (Nagai et al., 2016). The [^18^F]ASEM PET signal was observed in the same sites where PSAM4-positive neurons were found using immunohistochemical staining with FLAG, albeit sparsely (Fig. 2C-E). In addition, the [^18^F]ASEM PET signal was also observed in areas to which the PSAM4-5HT3-expressing regions project. This is most likely a visualization of PSAM4-expressing axon terminals, as we have successfully visualized axon-terminal expression of hM4Di in the SNr via DCZ-PET in a monkey whose putamen was injected with an AAV vector (Nagai et al., 2020). Although it remains to be confirmed by immunohistochemistry, [^18^F]ASEM PET can be used for *in vivo* anatomical mapping that provides valuable information for network analysis of chemogenetically induced activity change. Furthermore, [^18^F]ASEM-PET was also useful for delineating the brain regions of interest for subsequent functional verification. These results demonstrate the utility of PET imaging for validating expression of PSAMs in monkeys.

[^18^F]ASEM-PET failed to visualize PSAM4-GlyR expression (Fig. 5). This might not have resulted from low expression levels, because PSAM4-GlyR appears functional based on FDG-PET data (see below). Rather, the PET probe might have a lower *in vivo* affinity for PSAM4-GlyR, although a previous study showed that the same probe successfully identified PSAM4-GlyR in mice (Magnus et al., 2019). If this is the case, future work will be needed to develop more specific and higher affinity PET ligands for PSAM4-GlyR.

### Chemogenetic function of PSAM4-5HT3 in monkeys

In the present study, FDG-PET revealed that uPSEM817 was more effective than uPSEM792 in triggering neuronal activation via PSAM4-5HT3 (Fig. 4). In addition, phMRI studies showed that BOLD activity increased rapidly (∼13 min) in brain regions expressing PSAM4-5HT3 and persisted for at least 60 min following uPSEM817 administration (Fig. 6). These results are consistent with previous observations in mice that uPSEM817 has a lower effective dose than uPSEM792 (Magnus et al., 2019), and that its concentration in cerebrospinal fluid increases relatively rapidly and remains stable for more than 3 h after systemic administration in monkeys (Raper et al., 2022).

We have previously tested the DREADD excitability of striatal neurons in combination with hM3Dq and DCZ using FDG uptake as a marker and found that 1 μg/kg of DCZ increased striatal uptake by ∼10% (Nagai et al., 2020; Kimura et al., 2023), a similar range to what we observed for PSAM4 with 100 μg/kg of PSEM in the current study. Assuming comparable expression levels because the same AAV vector was used at similar titers in these cases, DREADD/DCZ may have greater efficacy in activating striatal neurons, although generalization by direct comparison is still limited. The latency of neuronal changes observed with PSAM/uPSEM817 in the phMRI study (13 min) was also relatively longer than what we have observed with hM3Dq-DREADD/DCZ (∼4 min; unpublished data), which may reflect the reported differences in pharmacokinetics that these ligands have in monkeys (Nagai et al., 2020; Raper et al., 2022). The dose-dependent effects and timing of activation suggest that behavioral experiments using PSAMs are feasible. If faster manipulation is desired, such as in electrophysiological studies, this may be possible by selecting a ligand with enhanced brain permeability relative to uPSEM817; for example, the use of uPSEM793 (the lowest effective dose in mice; Magnus et al., 2019) may be a candidate for this purpose.

### Limitations

First, the present study was primarily designed to validate the excitatory PSAM/PSEM system in primates, rather than the inhibitory PSAM4-GlyR system. Specifically, PSAM4-GlyR expressed in striatal neurons enhanced FDG uptake after administration, likely reflecting an excitatory chemogenetic effect, as previously reported (Gantz et al., 2021). Future studies should validate the inhibitory chemogenetic effect of PSAM4-GlyR on neuronal activity by injecting different brain regions and examining behavioral changes in NHPs, for which uPSEM817 should be the first choice.

Second, one of the most important technical issues in chemogenetics is ensuring that agonist ligands do not cause off-target effects on brain activity or behavior. For the DREADDs ligand DCZ, several laboratories including us have evaluated its effects on brain activity (Nagai et al., 2020; Fujimoto et al., 2022; Cushnie et al., 2023) and behavior (Nagai et al., 2020; Upright and Baxter, 2020) in non-DREADD-expressing monkeys and determined the upper dose limit for avoiding unwanted side effects. A previous study showed that administration of uPSEM817 (0.1 mg/kg) had no significant effect on heart rate or sleep in non-PSAM4-expressing monkeys (Raper et al., 2022). In the present study, we did not test animals that did not express PSAM. However, in phMRI experiments using a PSAM4-positive monkey, albeit under anesthesia, we confirmed that uPSEM817 administration (300 μg/kg) did not significantly affect brain activity in several control regions that did not express PSAM4. Future applications of the PSAM/PSEM system in monkeys would benefit from verifying the effects of actuators (PSEMs) on whole brain activity under awake conditions via multimodal imaging alongside behavioral testing in the absence of PSAMs.

Third, although we successfully visualized PSAM4-5HT3 expression in vivo using [^18^F]ASEM-PET, our *in vitro* confirmation via immunostaining with fusion proteins was limited (FLAG) or failed (HA). In our construct, the tag protein is fused to the N-terminals. It is possible that this construct is inappropriate and led to the failed immunostaining confirmation. Further validation is necessary for a postmortem reporter system using a fusion protein attached to the C-terminals or a co-expression system mediated by IRES or 2A sequences (cf. Magnus et al., 2019), or in situ hybridization assay. Additionally, comparing the PET reporter system with immunohistological labeling would allow quantitative accuracy of *in vivo* PET measurements to be determined.

Finally, while our phMRI data demonstrated that a sustained chemogenetic effect lasts for at least 60 min, the exact duration remains to be determined. This is important information for chronic NHP experiments and therefore needs to be addressed in a future study.

### Potential benefit of using the PSAM/PSEM system in NHPs

Given that DREADDs have already been shown to be effective in NHPs, the availability of PSAM/PSEM in monkeys will allow for additional validation with chemogenetics that act through channels, which is a simpler principle of neuromodulation than G protein-coupled receptors. Furthermore, in combination with the DREADD system, the PSAM/PSEM system allows multiplexing of chemogenetics; in combination with PET, visualization of brain circuitry could also be multiplexed. The ability to visualize the terminal region will facilitate pathway-selective manipulation by local agonist infusion (Oyama et al., 2021), and thus visualization and manipulation of multiple brain circuits in a single individual can be realized.

### Conclusions

Our multimodal imaging demonstration of neuronal modification by the PSAM/PSEM system illustrates the efficacy of these tools in monkey research. The information gained from this study— the optimal agonist, dose, and timing of chemogenetic neuronal manipulation via PSAM4—will facilitate the application of these tools in NHPs. Given the proven utility of the muscarinic DREADD system, this additional chemogenetic tool would increase the potential for flexible and reliable NHP circuit manipulation in a variety of situations.

## Acknowledgments

This study was supported by MEXT/JSPS KAKENHI Grant Numbers JP22H05157 (to KI) and JP20H05955 (to TM), by AMED Grant Numbers JP18dm0307007 (to TH), JP21dm0207077 (to MT), and JP16dm0107146 (to TM), by the cooperative research program (2021-A-14) at the PRI, Kyoto Univ, by JST (Moonshot R&D – MILLENNIA Program) Grant Number JPMJMS2295 (to TM and KI), and by the commissioned research by the National Institute of Information and Communications Technology (NICT; #22301), Japan (to TM). A monkey used in this study was provided by the National Bio-Resource Project: “Japanese Monkeys” of MEXT, Japan. We thank M.A.G. Eldridge for valuable comments on an earlier version of manuscript and J. Kamei, R. Yamaguchi, Y. Matsuda, Y. Sugii, T. Okauchi, T. Kokufuta, R. Yoshida, K. Yamashita, M. Nakano, M. Fujiwara, S. Shibata, and N. Nitta for their technical assistance.

